# Single molecule tracking the uncoupling of assembly and membrane insertion in Perfringolysin O

**DOI:** 10.1101/2021.05.26.445776

**Authors:** Michael J T Senior, Carina Monico, Eve E Weatherill, Robert J Gilbert, Alejandro P Heuck, Mark I Wallace

## Abstract

We exploit single-molecule tracking and optical single channel recording in droplet interface bilayers to resolve the assembly pathway and pore-formation of the archetypical cholesterol-dependent cytolysin nanopore, Perfringolysin O. We follow the stoichiometry and diffusion of Perfringolysin O complexes during assembly with 60 millisecond temporal resolution and 20 nanometre spatial precision. Our results suggest individual nascent complexes can insert into the lipid membrane where they continue active assembly. Overall, these data support a model of stepwise irreversible assembly dominated by monomer addition, but with infrequent assembly from larger partial complexes.

## Introduction

The Membrane Attack Complex Perforin / Cholesterol Dependent Cytolysin (MACPF/CDC) protein superfamily form giant multimeric *β*-barrel nanopores in the membranes of target cells. MACPF and CDC pores share a similar function, albeit on opposite sides of the battleground: MACPF proteins are a key component of the human immune system, using pore formation to lyse the cells of invading bacteria;^1^ whereas CDCs perform the analogous role in bacteria, and are virulence factors in a wide range of pervasive diseases including pneumococcal meningitis and listeriosis.^2,3^ Beyond just pore formation, MACPF/CDCs are also thought to play a role in processes as varied as histone modification,^4^ lipid raft aggregation,^5^ and the control of embryonic development.^6^ MACPFs and CDCs are linked by their shared *β*-barrel structural motif; ^7^ suggesting not only interesting evolutionary links between the two families, ^8^ but also strong similarities in the mechanism of pore formation. Thus understanding the assembly pathway for one branch of proteins has broad implications across this important superfamily.

The overarching model for MACPF/CDC pore formation is one where soluble monomers first bind to a cell membrane, then oligomerize into arc and then ring-shaped ‘prepore’ complexes, before finally folding cooperatively to insert a *β*-barrel pore that punctures the membrane.^7,9–16^ Detailed evidence supports this model, arising predominantly from a combination of protein structures and ensemble kinetics.^17–24^ Overall, this model focuses on the role of fully-assembled prepore and pore rings in membrane lysis; however beyond this well-established general scheme, much is still unclear regarding the kinetics of pore insertion.

To shed further light on pore assembly, we need better temporal and spatial resolution of the mechanism at a molecular level – where both pore dynamics and the nature of the forming complex can be resolved. Such experiments have the potential to help us evaluate the relative importance of the different factors that might control oligomerisation kinetics; to determine if *β*-hairpin formation by monomers is concerted; and to understand the precise interplay between membrane insertion and oligomerisation.

This approach has been pioneered by elegant single-molecule Förster Resonance Energy Transfer measurements of ClyA assembly kinetics in detergent micelles^25^ and by Atomic Force Microscopy (AFM). High-speed AFM has been used to examine the assembly of suilysin, ^15,26^ perforin, ^27^ listeriolysin,^14,28^ the membrane attack complex,^29^ and perforin-2. ^23^ Here we seek to improve on the temporal resolution rendered by AFM using single-molecule fluorescence imaging to track individual PFO pores as they assemble and insert into lipid bilayers.

PFO provides a well-studied archetype for CDC pore formation, ^30,31^ responsible for pore-formation in gas gangrene and necrohemorrhagic enteritis. Like other CDCs, PFO binds to lipid membranes in a cholesterol-dependent manner. Subsequent oligomerisation then leads to the formation of large (25-30 nm) pores that result in cell lysis.^21,22,31,32^ Here, using single-molecule fluorescence microscopy and optical Single Channel Recording (oSCR) in Droplet Interface Bilayer (DIB) model membranes^33,34^ we have tracked and quantified the stoichiometry of diffusing PFO complexes during assembly with 60 ms temporal resolution and 20 nm spatial precision.

We observe the membrane-associated growth and insertion of 1000’s of individual PFO pores. Overall our data showed that oligomers can continue growing after insertion, as judged by the reduction in diffusivity of the oligomer that is associated with its insertion into the membrane. In a recent preprint, Böeking and co-workers use a single-molecule dye release assay to link stoichiometry to pore formation for PFO on surface-tethered large unilamellar vesicles;^35^ their results also suggest assembly independent of membrane permeabilisation. Together, these results support a model where PFO arcs, currently thought to be kinetically trapped, can continue to grow post insertion.

### Exerimental design

To image PFO we took advantage of our previously developed DIB platform^34^ to track the assembly of individual monomers into large complexes and conductive pores. DIBs were created by the contact of two monolayers formed at oil-water interfaces on a sub-micron thick agarose film, allowing the bilayer to be imaged with single-molecule total internal reflection fluorescence (TIRF) microscopy. DIBs also enable control of the membrane potential, with GΩ electrical seals compatible with single-channel recording. ^36^

Bilayers were formed from mixtures of 1,2-diphytanoyl-*sn*-glycero-3-phosphocholine (DPhPC) and cholesterol. After bilayer formation, PFO was injected into the aqueous droplet to a final concentration of 37 nM using a piezo-controlled nanopipette (Fig.1A). A mixture of PFO labelled with Alexa-488 (PFO-a488) and unlabelled PFO (1:5.5 molar ratio) was used to optimise signal amplitude and minimise fluorescence self-quenching. ^37^

**Figure 1:**
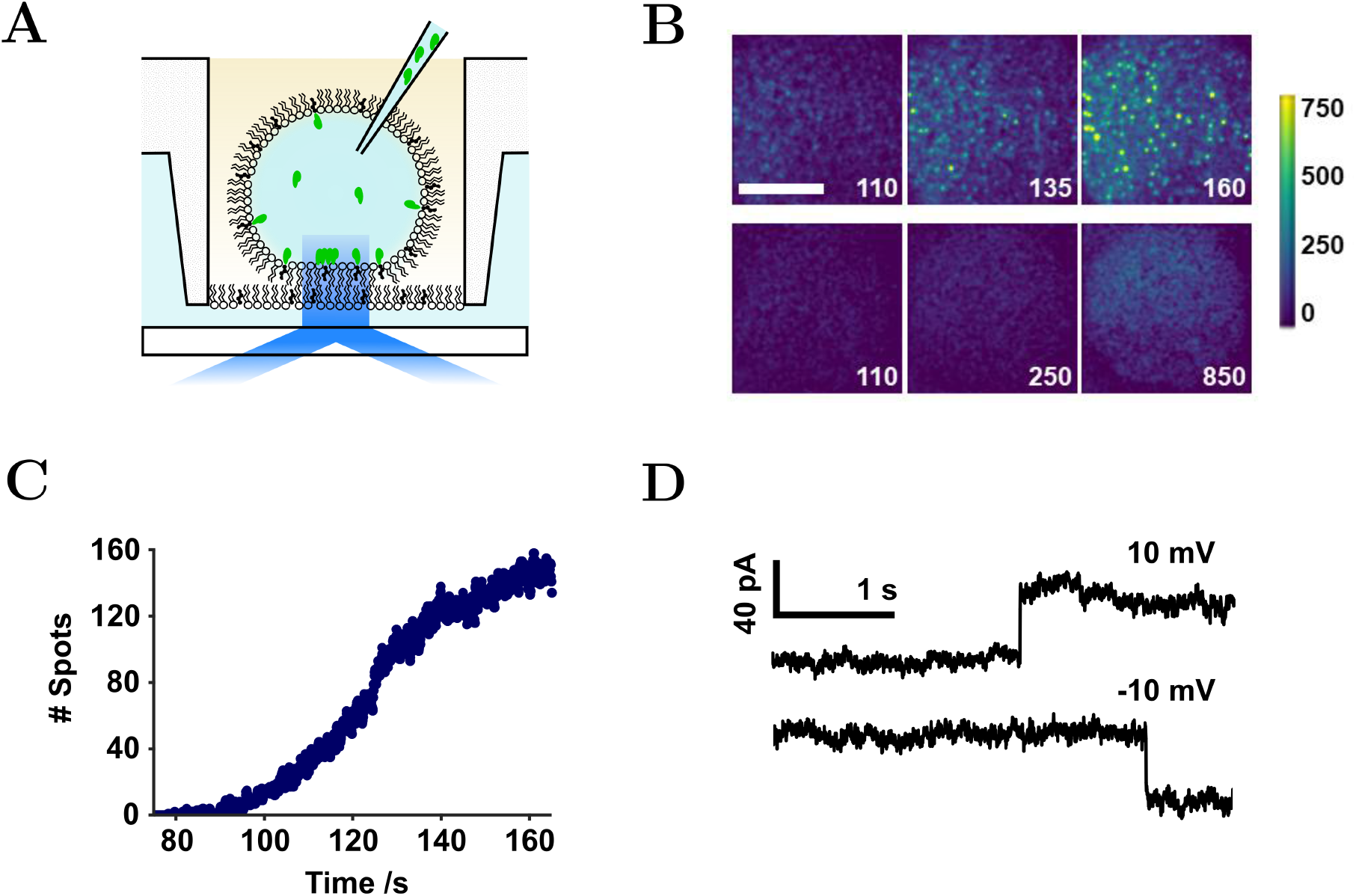
Experimental design. (A) Schematic of PFO (green) injection into a DIB contained in a microfabricated plastic well. The bilayer rests on a submicron thick agarose layer to enable TIRF imaging of the labelled protein on the bilayer. (B) Over time, PFO assembly was observed in bilayers containing cholesterol (upper) but not without (lower). PFO complexes, containing many labelled monomers appear as bright spots (green), with higher intensity than the monomer (blue). Images are blue-green false-coloured to aid discrimination, with a scale normalized to the intensity of a monomer. White numbering denotes seconds following protein injection. Image acquisition rate 16.67 Hz. Scale bar 25 μm. (C) Total number of spots detected in the video represented in B. (D) Electrodes placed in the hydrogel and the droplet permit measurement of current across the bilayer. Rarely, stepwise 4 nS changes in conductance were observed, consistent with insertion of single PFO pores into the bilayer.

Protein monomers were observed to bind to the bilayer and diffuse freely (Fig.1B). Analogous to our previous work on *Staphylococcus aureus α*-hemolysin, ^38^ spots with an intensity greater than that of a single fluorophore appeared over time (Fig.1B, Fig.1C, Movie S1) consistent with the formation of oligomeric PFO complexes. Trackmate, ^39^ a tracking algorithm implemented as a Fiji^40^ plugin, was applied to detect spots brighter than a single monomer (Fig.1C). Both the number and intensity of individual spots increased over time, before reaching a plateau (Fig.1C, Movie S1). PFO assembly was cholesterol dependent and binding was not observed in cholesterol-free bilayers (Fig.1B).

By inserting Ag/AgCl microelectrodes into the droplet and agarose substrate, the ionic flux across the bilayer was measured.^34^ Individual PFO insertion events were detected as rare stepwise changes in conductance of approximately 4 nS (Fig.1D), corresponding to previously reported values for PFO^41^ and other CDCs.^42–44^

### Tracking individual PFO complexes reveals a drop in diffusivity during assembly

The stoichiometries and time-dependent diffusion coefficients for complexes were determined. Fig.2 illustrates the assembly of four representative complexes from the dataset corresponding to Fig.1B. Each column in Fig.2 represents an individual complex. The position (Fig.2A), local pixel intensity (Fig.2B), time-dependent diffusion coefficient, and stoichiometry (Fig.2C) are shown. The time-dependent diffusion coefficients were calculated from the mean squared displacement over 3 second intervals. PFO stoichiometry was determined from the fraction of labelled PFO (1 a488 label per 6.5 PFO monomers) and normalisation of the time-dependent change in complex intensity to that measured for isolated single PFO-a488 monomers (Fig3). The stoichiometry increases essentially monotonically within an individual complex (Fig.2C). Higher stoichiometry complexes are also correlated with overall lower protein mobility (Fig.4B), with a dependence of the diffusion coefficient of a complex with its size.^45^

**Figure 2:**
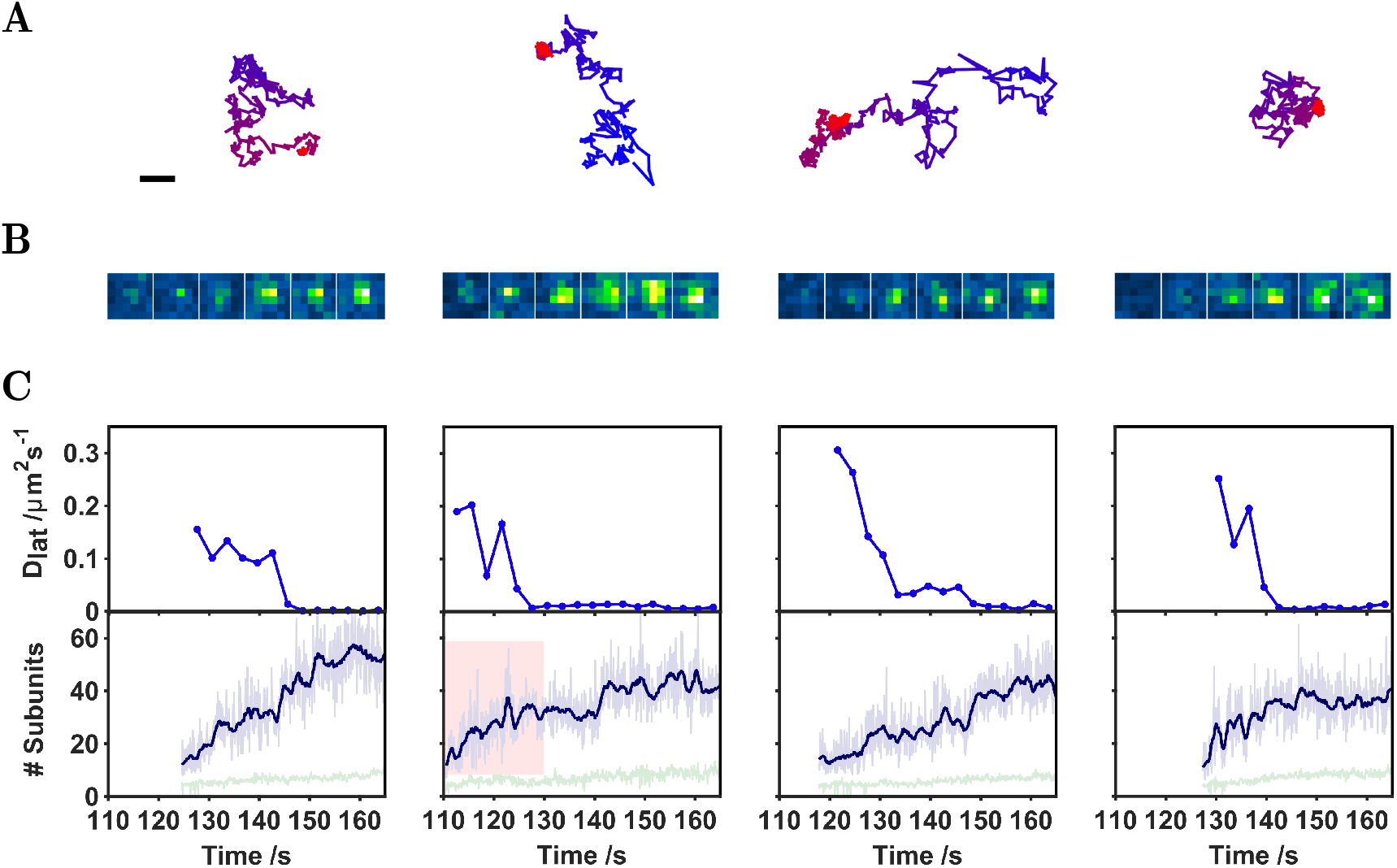
Assembly of individual complexes. Each column is an assembly event from our dataset, selected arbitrarily and representative of the majority of events. (A) Track coordinates of a complex over time (blue to red). Scale bar 1 μm. (B) Raw image intensity for a 6 × 6 pixel region of interest (ROI) coinciding with the appearance of the complex. Images are at 20 s intervals. (C) Time-dependent diffusion coefficient of the complex, calculated from the coordinates displayed in (A) (averaging over 3 s intervals); stoichiometry of the complex as a function of time. Raw values (light blue) and a 1.26 s running average (dark blue) is shown. The local background signal is also plotted (green). The red highlighted section corresponds to data used in Fig.3F.

Notably, complexes also show a marked drop in diffusion coefficient during assembly (Fig.2C). These sudden decreases in the diffusion constant occur independently of increasing stoichiometry, with continued complex growth following a decrease in diffusion coefficient (Fig.2C). We examined the relationship between stoichiometry and step-change in diffusion coefficient (Fig.4B). However this data does not reveal distinct populations associating diffusion with stoichiometry. We elected not to over-interpret the data. Further experiments may well reveal a more subtle relationship.

We expect the number of labelled monomers in a complex to follow a binomial distribution, so in a complex of 35 subunits a mean and standard deviation of 5.4 ± 2.1 labelled monomers is expected. The mean number of labelled subunits will increase linearly with the total number of subunits. The shape of the binomial distribution will affect the variance associated with the number of labelled monomers. At our largest complex size (approx 40), we estimate we estimate 6.1 ± 2.3.

### Mapping the distribution of single-molecule kinetics is consistent with biphasic, irreversible assembly

A key advantage of tracking individual complexes is the ability to examine the protein population without the need for extrinsic synchronisation of the ensemble (e.g., via chemical modification such as disulphide locking^20^). To analyse this collective behaviour, we examined all tracks that start with an initial stoichiometry below 20 subunits (to ensure effective averaging of the kinetics associated with early assembly). Trajectories were analysed by aligning the times at which each complex was first detected. Fig.4A depicts individual trajectories (light blue) and their median (dark blue). Overall, the assembly is described by an initial period of rapid growth, before a slower extension of the complex; best fit by a double-exponential (Fig.4A, red line). The corresponding pseudo-first order rate constants for assembly are *k*_1_ = 2.2 s^−1^ and *k*_2_ = 0.05 s^−1^. Of the two, *k*_2_ dominates (78%) and broadly matches the overall timescale for the number of complexes appearing over time (Fig.1C). The rate of assembly taken from this curve represents the number of successful collisions with monomers for a single complex per second. At the inflexion point when complexes are appearing most rapidly (125 seconds after injection in Fig.1C), the density of complexes is 0.03 μm^−2^. Thus, an estimate of the overall maximum rate for the assembly (rate of successful collisions between monomers and complexes) at this surface density is 0.0015 μm^−2^s^−1^.

Closer analysis of the rate of collision (*ν*) between monomers and nascent complexes can give further insight into the mechanism of complex assembly. We thus estimated collision rates using a simple 2D kinetic model of collision and diffusion:^50^ Making the assumption that complexes can be approximated as static sinks relative to monomers, and that collisions occur below a fixed encounter radius,

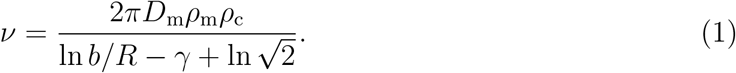

Here *ρ*_c_ = 0.03 μm^−2^ is the surface density of complexes determined from the number of complexes per area at the inflexion point in Fig.1C; *ρ*_m_ = 14 μm^−2^ is the surface density of single free monomers; estimated by dividing the overall intensity per unit area of the bilayer at the inflexion point by the intensity of a single monomer (Fig.3A,B). The diffusion coefficient for single monomers (*D*_m_ = 1.5 μm^2^ s^−1^) was determined from the mean square displacement vs. time for labelled monomers diffusing on the bilayer. The encounter radius within which a collision is deemed to have occurred, *R*, was set at 2.75 nm, approximating the radius of a PFO monomer.^17^ *γ* = 0.577 the Euler-Mascheroni constant and 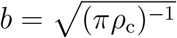. The rate for monomer-complex collisions determined from this analysis is *k* = 0.58 μm^−2^ s^−1^. Comparing the rate for monomer-complex collisions to the overall rate of complex assembly (0.0015 μm^−2^ s^−1^), approximately 1 in 390 collisions between monomers and nascent complexes are successful and result in growth.

**Figure 3:**
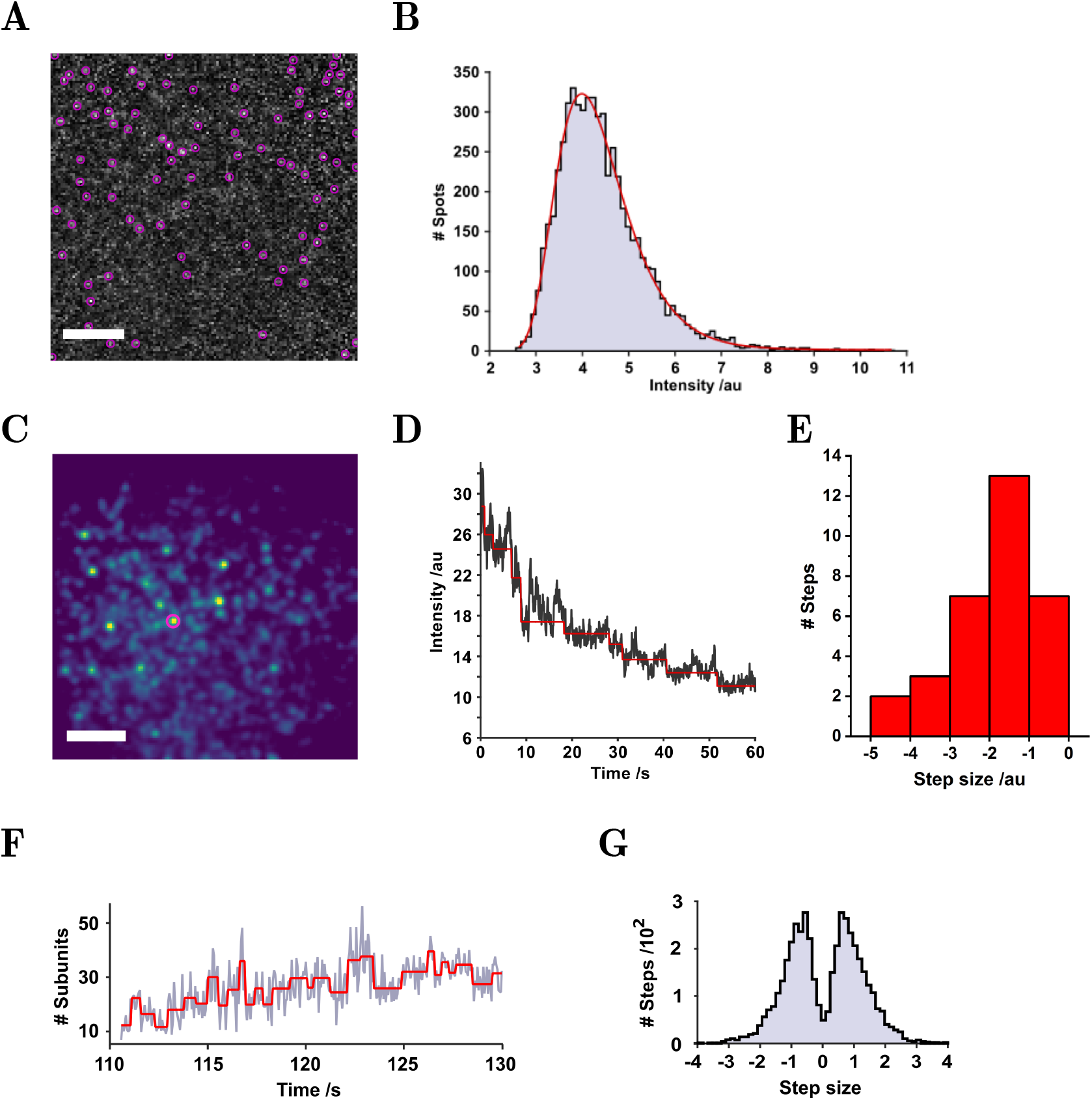
Intensity calibration of PFO-a488 stoichiometry. A&B): Estimating single-fluorophore intensity by tracking single labelled monomers. (A) PFO-a488 bound to a DIB membrane at a concentration (100 pM) insufficient to initiate assembly. Diffusing monomers were tracked as indicated by the purple circles. Scale bar 10 μm. (B) Following normalisation to the distribution of incident laser intensity, a histogram of pixel intensities from the detected monomer spots in a 50-frame window was plotted. The histogram was fitted with a log-normal function and the modal intensity used to calibrate stoichiometry during assembly. The intensity of each assembled complex is multiplied by the labelling ratio and divided by the singlefluorophore intensity, to estimate the number of monomers within an individual complex. (C-E): Estimating single-fluorophore intensity by photobleaching. Subunit counting by photobleaching ^46,47^ was undertaken with PFO-a488. Plotting the intensity of complexes over time and then applying a Chung-Kennedy stepfinding algorithm ^48^ to the traces highlighted the stepwise decrements due to photobleaching. (C) PFO-a488 complexes were assembled on a DIB in the dark, before illumination and imaging to photobleach individual complexes (0.9 mW at the objective back aperture). Scale bar 10 μm. (D) Mean intensity for the spot highlighted in (C). Red line is the result of Cheung-Kennedy step-finding filter. (E) Histogram of the step-sizes from photobleaching spots in D, median step-size = –1.4 a.u. After background correction and accounting for the ratio of labelled to unlabelled PFO the number of monomers per complex in this image was estimated at 32–65, in agreement with our intensity-based calibration. (F-G): Estimating monomer intensity using stepwise fluctuations within an individual, nascent PFO complex during assembly. (F) A step-finding algorithm ^49^ applied to a stoichiometry trajectory from Fig.2C (highlighted as a shaded red box) shows step-wise changes in intensity.(G) A histogram of step-sizes, normalised to the intensity of a single fluorophore, indicates the most common event is a single monomer arriving at or leaving the vicinity of the complex

This high propensity for successful assembly is very different to our previous work on the heptameric *β*-barrel toxin *α*-hemolysin, ^38^ where only 1 in 10^7^ collisions were successful. Whereas *α*-hemolysin requires reversible assembly, the high assembly probability for PFO provides evidence for a different, irreversible, assembly mechanism consistent with previous work.^15,51^ An irreversible stepwise monomer-addition mechanism for PFO is arguably expected simply based on the large number of monomers required to form MACPF/CDC pores.

Fig.4C depicts the variation in the distribution of PFO stoichiometries with time, aligned to synchronise the start of assembly for all complexes. Here, tracking parameters were optimised to now detect monomers as well as protein complexes. As expected, a shift from smaller to larger complexes with time is observed with a broad distribution of final complex sizes (30 – 40 subunits). The number of low-stoichiometry complexes and monomers in the first 5 seconds is high (19200) relative to the later peaks, due to the excess of monomers on the bilayer at early stages of assembly.

**Figure 4:**
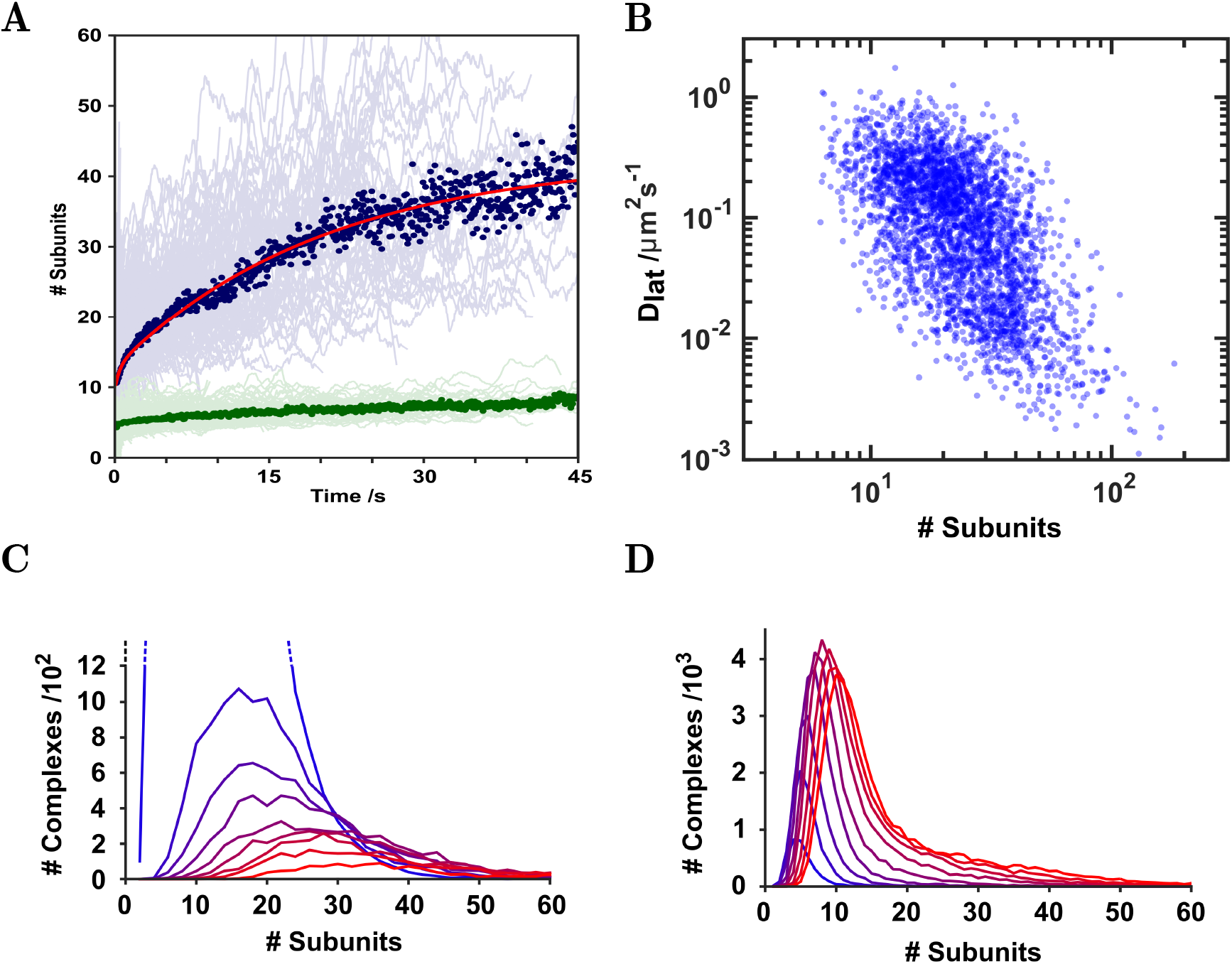
Overall kinetics of assembly. (A) Assembly events were aligned by start time and plotted (light blue lines). A threshold on trajectories was applied to select those with an initial stoichiometry of 20 subunits or fewer to ensure temporal alignment captured early assembly. The median of these trajectories is superposed (dark blue dots), representing the average rate of assembly of a single complex, best fit to a double exponential (red line). The equivalent background signal from these trajectories is also shown (green). (B) Trajectories for a representative 500 tracks were segmented at 1.2 second intervals and the diffusion coefficient determined for each segment. The diffusion coefficient was then plotted against mean stoichiometry for each corresponding segment. An approximately power law relation between *D*_lat_ and stoichometry was observed (*D*_lat_ ∝ *S*^−3.8^).(C) Tracking parameters were then adjusted to include monomers (as opposed to just complexes as per (A), tracks were aligned, and stoichiometry histograms corresponding to 5 second intervals are plotted and color-coded (blue to red). The histogram peak position shifts over time in the assembly process, as a large number of small oligomers (blue) are converted to smaller numbers of larger complexes (red). (D) Equivalent histograms as (C), but without alignment of the tracks shows the overall evolution of PFO during assembly. Histograms show the change in overall PFO complex size since the injection of protein into the droplet. Each histogram corresponds to a 10 second interval (color-coded blue to red).

We also examined the events in Fig. 4C not relative to assembly, but in ‘real-time’ – sampled every 10 seconds after injection of the protein in the bilayer (Fig.4D). The overall peak area increases over time as more monomers bind to the bilayer. In addition, over time the tail of this distribution elongates as the mean stoichometry transitions towards higher, assembled, complexes. However, there are always more monomers on the bilayer than complexes, and the low-stoichiometry peak dominates.

We examined the time-dependence in stoichiometry within an individual complex (Fig.4D). For these data, we observed significant fluctuations in intensity within the overall upward trend. To explore the relevance of these fluctuations we used a step-finding algorithm^49^ (Fig.3F). A histogram of the resultant step-sizes (Fig.3G) exhibits two peaks with a median step-size of ±1 monomer intensity units. This indicates that the majority of steps found by the algorithm correspond to a single labelled monomer in the vicinity of the complex. It is tempting, but we believe ultimately incorrect, to interpret these steps as individual monomers entering and leaving a complex: Given the frequency of successful collisions relative to the maximum stoichiometry of a final complex, it is unlikely that these frequent steps correspond to the dynamic addition or removal of monomers from a nascent complex. It is more likely that the dominant contributor to these observed steps is the diffusion of monomers nearby (i.e., within our ability to spatially resolve) the forming complex, rather than active assembly.

### Rare plasticity events are observed during PFO assembly

After characterising the overall trend of monomer addition and growth, we examined more closely those events that showed evidence of interaction between larger nascent complexes.

During assembly, diffusing multi-subunit complexes were observed to encounter each other and combine. As our spatial resolution is diffraction-limited, to correctly define a merging event those complexes must not only reside within the same spot, but also persist for longer than the expected diffusion time for a single complex. For comparison, we observed 30 merging events in more than 500 tracked complexes.

Fig.5A and Movie S2 present examples of plasticity events: the montages in Fig. 5A highlight the merging events as they appear for each event (at -10, -5, -2.5, -1.25, 0, +1.25, +2.5, +5 seconds relative to the event). The stoichiometry of the resulting complex closely matches the combined stoichiometry of the initial complexes. Fig.5B and Fig.5C are the corresponding histograms of PFO stoichiometry for before and after each merging event. The median complex stoichiometries before and after merging are 26 and 58 respectively. We also observe changes in diffusivity associated with plasticity: Events were observed in which a mobile complex joins another mobile complex; the two diffusing complexes then merge, exhibit a concomitant decrease in mobility, and then subsequently diffuse away as a single complex.

**Figure 5:**
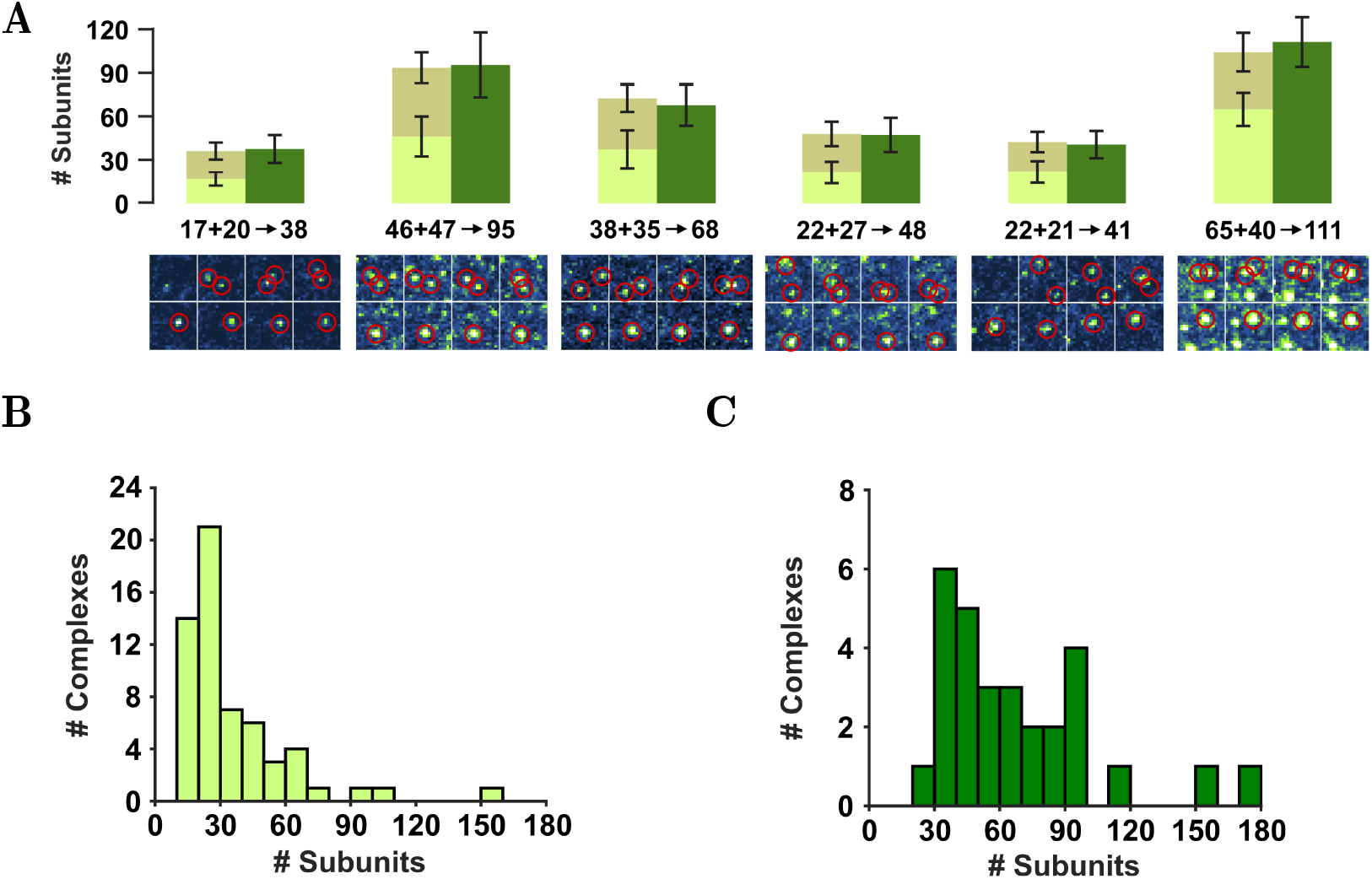
PFO plasticity. (A) Partial complexes we observed to join to form larger complexes; events were analysed for their stoichiometries, by averaging 50 frames before (light green/brown) and after (dark green) the merging event. The position of each complex is highlighted in the image sequence below, the frames displayed being taken at -10, -5, -2.5, -1.25, 0, 1.25, 2.5 and 5 seconds relative to the event. (B) Histogram of complex stoichiometries before merging (median = 26). (C) Histogram of complex stoichiometries after merging (median = 58).

### Pore formation appears infrequent relative to assembly

After assembly, PFO forms pores in the membrane. To follow this process we combined optical Single Channel Recording^36^ with single-molecule fluorescence tracking to enable comparison between pore formation and assembly. The presence of pores in the membrane was detected by imaging Fluo-8, a calcium sensitive dye, while the extent of oligomerization of monomers into complexes was measured by imaging PFO labelled with Alexa-647 (Fig.6A). Alexa-647 was chosen to enable simultaneous imaging of both calcium flux and oligomerisation. As expected, ^38,52^ the ionic flux-induced fluorescence intensity of a single pore varies linearly with the voltage applied across the bilayer (Fig.6D).

**Figure 6:**
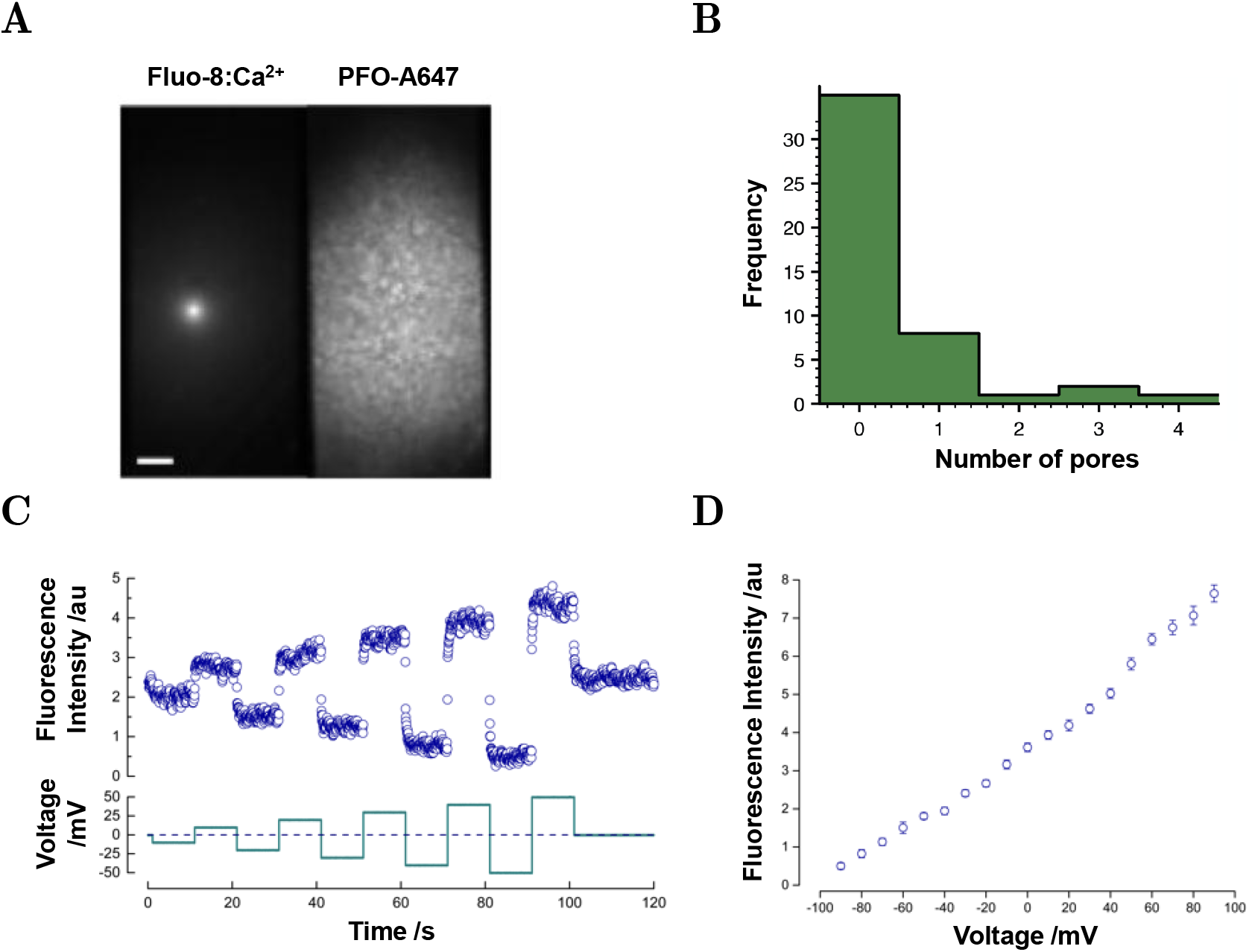
Optical single channel recording and dual-color imaging of pore formation. (A) Pores are detected optically in the left channel from fluorescence emitted by Fluo-8, a calcium sensitive dye as it flows through the pore. The right channel reports on fluorescence from PFO labelled with Alexa-647. Pores form rarely relative to the large number of assembled complexes. Scale bar 2μm. (B) Distribution of PFO pore events per bilayer as determined by Fluo-8 imaging of calcium flux. (C) Time dependent responses to applied steps in membrane potential from a single PFO pore present in a DIB, corresponding to (A). (D) Fluorescence-voltage response of a PFO pore. Error bars represent the standard deviation from the mean fluorescence intensity.

Surprisingly, relative to the number of assembled complexes on the bilayer, we only observed rare conductive pores (Fig.6B). Pores were only detected under conditions where individual PFO complexes could not be resolved by single-molecule fluorescence due to high concentration (Fig.6A, right channel). We are limited by the maximum stoichiometry of the assembled complex. If the concentration were increased from 37 to 100 nM by addition of unlabelled PFO, stoichiometry would decrease by factor of 2.5. Thus every pore would only have an average of 2 labels and would be unlikely to be detected over the background. We expect the complexes in Fig.6A (right) to be mainly assembled because concentration is 2.5 times higher than in Fig.4. These experiments were challenging because at low protein concentration no changes to conductivity were observed, but at high concentration bilayers ruptured suddenly, before any insertion events were observed or even before the electrode was inserted into the droplet, consistent with a cooperative nature of assembly. Nonetheless, simultaneous electrical and optical recordings show that once formed, PFO pores were extremely stable (often lasting several hours) with stable conductance (Fig.1D, Fig.6A,C).

## Discussion

Our measurements provide a broad overview of PFO assembly from early to late stages of pore formation (Fig.1 & Fig.2). We observe bi-component assembly kinetics (Fig.4) with rapid oligomerization in early stages followed by a period of slower extension of the oligomer. These kinetics are reminiscent of the two-step mechanism reported for streptolysin-O, where membrane-bound dimerisation is followed by sequential oligomerization, ^53^ and for the initial conformation change reported for PFO^54^ and other CDCs.^16,55^ A two-step process is also consistent with the observations of Böcking and co-workers. ^35^ In light of these reports it is tempting to assign our fast-process to dimerisation. However in this experiment, to ensure only assembling complexes are selected, our tracking parameters exclude signals from monomeric PFO. We therefore have insufficient discrimination to assign our fast kinetics to dimerisation. Indeed, examining Fig.4A closely, *k*_1_ dominates over low subunit numbers up to ≈ 15, much larger than just dimerisation. A second possibility is two assembly rates corresponding to pre- and post-insertion of the nascent complex.

We did not observe any evidence of PFO binding to the membrane as higher-order oligomers. In our intensity calibration, 100% labelled monomers have a single intensity state (Fig.3A,B) corresponding to a single step size both during assembly (Fig.3C-E) and photobleaching (Fig.3F,G). Using this calibration, the largest size of detected complexes corresponded to the expected size of a PFO pore^21,32^. Monomer intensity was measured at concentrations insufficient for assembly (100 pM). There would be a possibility of oligomers becoming dominant at higher concentrations (37 nM). However, the dominant mechanism we observed was monomeric stepwise growth (Fig.3G); if large oligomers were binding, we would have seen them as large steps in assembly. We did detect some complexes combining, but these are very rare (Fig.5).

Although there is a broad distribution of individual assembly rates, in essence all complexes follow the same assembly pathway (Fig.4A), with little evidence of different modes of assembly or trapped intermediates. The distribution of PFO stoichiometry over time (Fig.4C) is characterised by a broad distribution in the number of subunits throughout all stages of PFO assembly, consistent with the reported variable pore sizes typical of CDC oligomers.^13,15,21,56^ Additionally, Tweten and colleagues used NBD attached to a cysteine-substituted derivative of PFO to suggest that oligomerization is favoured over initiation of new complexes, ^41^ a model supported by our data.

Our results support a model of PFO assembly consistent with an irreversible stepwise addition of monomers to assembling complexes. This is mainly evidenced by the continuous growth to larger complexes (Fig.2, Fig.4) and by the number of successful collisions. Only 1 in 340 collisions lead to successful oligomerization, surprisingly frequent compared to reversible assembly of other oligomeric proteins.^38^ It’s possible that this difference may be accounted for by the cooperative nature of *α*-hemolysin assembly, whereas the scale of CDC oligomers may instead require some kind of switching mechanism to retain monomers once they join.^55^ This supports an irreversible model of CDC assembly as also evidenced by cryoEM and AFM.^15,51^

Examining the assembly of individual complexes, Fig.2 shows fluctuations in intensity are consistent with the addition of single, but not multiple, fluorescently labeled monomers. Given our labelling ratio, (1 fluorophore per 6.5 monomers), the occurrence and detection of oligomers smaller than this size would be low. Once complexes appeared they do not dissociate (Movie S1). This is also consistent with a model based on irreversible assembly of monomers and in agreement with previous reports,^18,51^ where completion of existing complexes is favoured over the initiation of new complexes. Ultimately, our insight into these early kinetics is limited by the the competing requirements of sub-stoichiometric labelling and the need to discriminate between nascent complexes and the surrounding milieu of monomers.

We were able to track the assembly of individual complexes with 20 nm precision, calculated from the error in fitting of a Gaussian point spread function to diffraction limited spots in our data. Closer examination of these tracks also revealed a significant drop in the diffusion coefficient during assembly (Fig.2). A likely cause for this change would be the transmembrane insertion of a portion of the complex which becomes trapped on the underlying agarose matrix. This would be consistent with previous reports of transmembrane insertion of oligomers prior to full pore formation. ^13,15^ Based on this interpretation, we see that oligomerisation continues following transmembrane insertion (Fig.2C); suggesting PFO arc insertion, the formation of a semi-toroidal pore, and then continued on-pathway growth of the complex towards a final state with a stoichometry consistent with a completed ring.

Overall, similar to Böking and co-workers, ^35^ our interpretation fits with the sequential insertion model for *β*-barrel formation,^24^ where the conformational change propagates along the oligomer. In this model, the semi-toroidal pore forms during assembly and presumably insertion proceeds along the partially-inserted oligomer.^57^ A model for pore formation by perfringolysin similar to that proposed here and in the accompanying paper ^35^ was previously put forward for the CDC streptolysin, firstly on the basis of the imaging of pores in membranes;^58^ later investigated further by tracking the appearance of growing oligomers ^53^ and via further imaging studies with the benefit of a mutant protein modulating the size of membrane pores formed.^59^ That model envisaged dimerisation on the membrane as the initial step in pore assembly^35,53^ and that the early membrane lesion is lined by a free edge of the lipid membrane and extended gradually during oligomerization. ^59^ Subsequent studies appeared to discount this model^41^ for pore formation. However, the real-time analysis of large numbers of oligomers forming and of pore formation (this paper, ^35^) suggests that the insights gained previous from studying streptolysin may indeed have been correct after all.

While post-insertion arc growth needs further investigation to determine whether different CDCs and/or different experimental systems lead to different outcomes, it is nevertheless tempting to speculate how membrane-bound PFO monomers in the prepore state can join an arc that has undergone a large conformational change to insert into the membrane. One hypothesis is that the PFO monomer may interact with the inserted arc, followed by vertical collapse and unfurling of the *β*-hairpin. This model is akin to the sequential insertion mechanism. An alternative hypothesis is that PFO monomers may undergo a conformational change as a result of interacting with the toroidal lipid edge and subsequently add to the arc pore.

We were surprised that our optical Single Channel Recording (Fig.6) showed that only few of these inserted complexes were able to conduct ions. CDC pore formation kinetics is temperature dependent,^41^ however, our efforts to repeat these experiments at higher temperature (37 °C) resulted in no appreciable increase in kinetics of pore formation. PFO insertion has also been reported in liposomes prepared with similar lipid mixtures of DPhPC and Cholesterol. ^60^ In isolation, these data would support an interpretation where transmembrane membrane insertion of one or more *β*-hairpins occurs, but not formation of a conductive pore. Interestingly, these observations are significantly different to those reported by Bocking and co-workers, ^35^ where membrane permeabilisation of PFO on large unilamellar vesicles was a predominant feature. Indeed, studies of PFO and other pore-forming toxins demonstrate greater activity of conductive pores in LUVs^20,61^, supported lipid bilayers^15^ and planar bilayers^41^ than we report here in DIBs. Further experiments are needed to ascertain the cause of this difference. However, our result here is in contrast to our work on other pore-forming toxins, where pore formation is commonplace. ^38,62^ To speculate further, the membrane tension and curvature are two obvious physical parameters that differ between SUV and DIB membranes, and would be expected to affect permeabilisation; particularly in the case of semi-toroidal pores.

Fusion of incomplete arc-shaped complexes has been hypothesised as a mechanism of pore assembly,^58,63,64^ supported by time-lapse AFM images for listeriolysin O (5 minutes time interval, for a total of 4 consecutive images).^14,28^ Additionally, disassembly and reassembly of such structures was also observed in PFO.^43^ Our single-molecule data show real-time evidence of plasticity in PFO complexes (Fig.5), supporting a model in which the fusion of partially assembled oligomers contributes to pore formation in CDCs. Although clearly present, these events represent a relatively small fraction (≈ 0.1%) of the overall assembly process which must be dominated by monomer or small (*n <* 6) oligomer addition.

In summary, single-molecule fluorescence experiments provide a higher time-resolution complement to the higher spatial resolution single-molecule information offered by AFM and EM. The ability to follow in real-time every single CDC complex without the need for chemical synchronisation of the population has brought new information to improve our understanding of how these important protein pores assemble. Our results suggest the insertion of incomplete arc-shaped membrane lesions might be a common feature of the CDC pore formation mechanism, distinct from the canonical model of prepore assembly prior to insertion.

## Methods and Materials

### PFO

A single cysteine mutant of PFO (PFO[C459A, E167C]) and PFO[C459A] were expressed in *E. coli* and purified.^65^ PFO[C459A, E167C] was 1:1 labelled via maleimide coupling chemistry with Alexa-488 or Alexa-647. Deletion of the native Cys residue has previously been shown to have an impact on the haemolytic activity of the protein^37^ but the difference in activity would only be apparent at concentrations < 1 nM, well below those used in this study. E167 was chosen as it is situated on the ‘top’ of the monomer, relative to the membrane, and so does not influence the pore assembly mechanism.^66^

Alexa-labelled PFO was separated from free dye by gel filtration in 50 mM HEPES, 100 mM NaCl, pH 7.5 and stored with 10% glycerol at -80 °C. Labelling efficiency was 100% for Alexa-488 and 68% for Alexa 647. A haemolytic assay was performed to verify the biological functionality of the protein. PFO was reconstituted at pH 6.5^60^ in fluorescence imaging buffer (0.5 M KCl, 10 mM MES, 0.1 mM EDTA, 1 mM Trolox and 10 mM Cysteamine) before injection into the droplet.

### Droplet Interface Bilayers

DIBs were created by the contact of two monolayers formed at oil-water interfaces. First, a monolayer was formed around a 138 nl aqueous droplet with 8.5 mg ml^−1^ of 1,2-diphytanoyl-*sn*-glycero-3-phosphocholine (DPhPC) in 10% silicone oil in hexadecane. Meanwhile, a second monolayer of 15.5 mg ml^−1^ 1:2 DPhPC:cholesterol in hexadecane was formed on a sub-micron thick agarose film (0.75% w/v low-melt agarose in water), spin-coated on a coverslip within a PMMA micro-machined device.^34^ 1.8% w:v of low melt agarose in the same buffer, was added around the PMMA chamber for hydration. The monolayers were incubated for 2.5 hours at RT, and then the 138 nl droplet was pipetted into the well to form a bilayer upon the contact of the two monolayers. The bilayer was then heated for 30 minutes at 37°C to allow equilibration of the asymmetric monolayers (this is longer than the half-life for cholesterol flip-flop (50 min at 25°C^67^)). Subsequently the agarose film was further rehydrated by fusing a 2.3 nl volume of buffer to it from a nanolitre piezo-injector (World Precision Instruments). Once the imaging conditions had been optimized, 4.6 nl of PFO (PFO-a488:PFO, 1:5.5) in solution was piezo-injected into the 138 nl droplet to a final total concentration of PFO in the droplet of 37 nM. Imaging was initiated not more than 30 seconds after injection of the protein.

### Imaging and data analysis

Total Internal Reflection Fluorescence imaging of the bilayer following injection of the protein solution was performed using a customized Nikon Eclipse TiE inverted microscope equipped with a *×*60 1.45 NA oil-immersion objective. Videos were recorded on an electron-multiplying camera (Andor Ixon). Events were tracked using the TrackMate plugin for Fiji/ImageJ. ^39,40^ Once the coordinates had been extracted for each spot in the tracks, a Gaussian function was fitted to the spots at the specified coordinates in the original video, to extract the spot amplitude, background and x, y positions. This fitting and all further analysis was performed using custom software written in Matlab. Diffusion coefficients for 3 second (Fig.2) and 1.2 second (Fig.4A) intervals were determined by fitting of the mean-squared-displacement vs time.

### Electrical recordings and dual-colour imaging

138 nL droplets contained Fluo-8 20 ug mL^−1^ in 10 mM MES buffer pH 6.5, 1 M KCl, 0.14 mM EDTA, and hydration agarose (2% w:v) in 0.5 M CaCl_2_, 0.1 mM EDTA in 10 mM MES pH 6.5. A symmetric bilayer was prepared with 16 mg ml^−1^ of DPhPC:Chol 1:2 in 20% silicone oil in hexadecane. A ground electrode was added to the agarose and the reference electrode to the droplet connected to a CV-203BU headstage and an Axopatch 200B amplifier (Axon Instruments). The droplets and electrodes were contained in a Faraday cage. An emission image splitter (Cairn OptoSplit) was added in the camera to enable simultaneous recording of the green (Fluo-8) and FarRed (PFO-a647) channels. Initially, a mixture of 1:5.5 PFO-a647:PFO was injected to the droplet to a final concentration of 45 nM, and equivalent assembly videos were acquired, as in the case of 1-colour imaging with PFO-a488. For experiments in Fig.5 the PFO concentration was increased to 100 nM.

## Abbreviations

CDC: (Cholesterol-Dependant Cytolysin)
PFO: (Perfringolysin O)
MACPF: (Membrane Attack Complex)
DIB: (Droplet Interface Bilayer)
AFM: (Atomic Force Mi-croscopy)
TIRF: (Total Internal Reflection)
oSCR: (Optical Single-Channel Recording)

## Declaration

The authors declare no conflict of interest.

## Acknowledgement

MJS was funded by the EPSRC Life-Science Interface Doctoral Training Centre at Oxford University. We thank Till Böcking and James Walsh for their helpful comments during the preparation of this manuscript.

## Supporting Information Available

Supporting Figures and Movies.

